# Strontium treatment potentiates bone anabolic action of intermittent PTH in ovariectomized rats

**DOI:** 10.1101/2025.09.18.677152

**Authors:** Cyril Thouverey, Isabelle Badoud, Patrick Ammann

## Abstract

To optimize osteoporosis therapy with the parathyroid hormone fragment teriparatide (PTH1-34), we sought to determine whether strontium (Sr) could potentiate the bone anabolic action of intermittent PTH1-34 in ovariectomized rats. Female rats were either Sham-operated or ovariectomized (Ovx) at 6 months of age. Eight weeks after surgery, Ovx rats received either vehicle solutions, 625 mg/kg/day Sr (5 days per week), 8 µg/kg/day PTH1-34 (5 days per week), or the combined treatments for 8 weeks. PTH1-34 reversed Ovx-induced deterioration of trabecular microarchitecture, apparent volumetric bone mineral density (vBMD), and strength, whereas Sr alone increased tissue-level vBMD without significantly affecting trabecular bone mass. Co-treatment with Sr and PTH1-34 further increased trabecular thickness, apparent and tissue-level vBMD, bone material properties (force and working energy), and trabecular bone strength compared with PTH1-34 alone. In cortical bone, PTH1-34 increased bone volume, cortical thickness, and apparent vBMD, while co-treatment further enhanced cortical thickness and apparent vBMD, maintained the Sr-induced increase in tissue-level vBMD, and significantly improved bone strength. In primary osteoblast cultures, Sr and PTH1-34, administered either alone or in combination, increased *Rankl* and decreased *Opg* expression, consistent with the elevated urinary levels of the bone resorption marker deoxypyridinoline in vivo. Sr or PTH1-34 alone stimulated *Igf1* and *Alpl* expression, whereas co-stimulation further enhanced these osteogenic markers. In conclusion, combining Sr with PTH1-34 integrates the osteoanabolic effects of PTH1-34 on bone mass with the mineral-level effects of Sr on bone material properties, leading to synergistic stimulation of bone formation and superior improvements in bone quality and strength.

## Introduction

A large arsenal of approved therapies is currently available to treat osteoporosis. There are anti-resorptive drugs such as denosumab (antibody neutralizing receptor activator of nuclear factor-κB ligand antibody -RANKL-) and bisphosphonates that inhibit bone resorption, and osteoanabolic agents such as romozosumab (sclerostin-neutralizing antibody), teriparatide (first 34 amino-acids of parathyroid hormone -PTH1-34-) and abaloparatide (parathyroid hormone-related peptide analog) that stimulate bone formation. Each of these anti-osteoporotic treatments has its own strengths and weaknesses in use [1]. Therefore, a significant part of research is currently dedicated to the optimization of therapeutic options for osteoporosis, i.e., the combination or the sequential use of different drugs [1].

Daily injections of PTH1-34 (intermittent PTH1-34 treatment) increase bone mass and reduce vertebral and non-vertebral fracture risk in osteoporotic patients [2, 3]. The bone anabolic action of PTH1-34 is attributed to its ability to increase osteoblast differentiation and activity, resulting in accelerated deposition and mineralization of new bone matrix [4, 5]. Those effects involve direct action of PTH1-34 on osteoblasts [6] or indirect actions through decreased secretion of the bone formation inhibitor sclerostin by osteocytes [7, 8] and increased local production of growth factors such as insulin-like growth factor 1 (IGF1) by osteoblasts [9, 10]. PTH1-34 also enhances bone resorption by inducing RANKL secretion by osteoblasts and osteocytes, but since bone resorption and formation are coupled, PTH1-34-induced increase in bone formation creates a positive balance in each bone remodeling unit affecting the skeleton [4, 5]. However, the accelerated bone remodeling induced by intermittent PTH1-34 treatment may limit improvements in intrinsic bone tissue properties, including collagen maturation and mineralization degree, thereby potentially constraining gains in bone material quality despite increases in bone mass [11]. PTH1-34 therapy is only used for severe cases of osteoporosis and recommended for a limited period of two years because of potential side effects such as osteosarcoma and hypercalcemia observed in preclinical studies [12].

Strontium (Sr), in the form of strontium ranelate, is no longer distributed as a pharmacological therapy for osteoporosis because of potential cardiovascular events but is still commercialized as a bone health supplement. Sr treatment reduces vertebral and hip fracture risk in postmenopausal women [13, 14]. Mechanistically, Sr has demonstrated a dual mode of action in clinical and preclinical investigations by simultaneously inhibiting bone resorption and stimulating bone formation [13, 15]. In vitro, Sr restricts osteoclast development and resorbing activity either directly or by decreasing the *Rankl*/*Opg* expression ratio in osteoblasts [16, 17]. In addition, Sr stimulates osteoblast proliferation and differentiation [17, 18]. Despite these effects, Sr monotherapy generally induces only modest increases in bone mass and architecture in preclinical models, suggesting that its primary contribution may reside at the material rather than structural level of bone [19, 20]. At the tissue level, Sr can also substitute calcium in bone hydroxyapatite and improve bone material properties [19, 20].

Combining PTH1-34 and Sr therapies represents a rational strategy to enhance treatment efficacy by integrating the strong osteoanabolic effects of PTH1-34 on bone mass with the mineral-level effects of Sr on bone material properties, thereby addressing complementary limitations of each monotherapy. Since the two therapies act through distinct but partially overlapping mechanisms in bone, we hypothesized that combination of PTH1-34 and Sr treatments could synergistically augment bone mass and strength in a rat model of postmenopausal osteoporosis.

## Materials and Methods

### Study design

Seventy 6-month-old Sprague-Dawley female rats (Charles River Laboratories, L’Arbresle, France) were housed individually at 25 °C with a 12/12-hour light/dark cycle. The rats were strictly pair-fed a laboratory diet containing 15% casein, 0.8% phosphorus, 1% calcium, 70–80% carbohydrates, and 5% fat (Kliba Nafag, Kaiseraugst, Switzerland) to the Sham-operated group to ensure identical caloric intake across groups and were randomly assigned to 5 groups of 14 animals. Four groups of rats were ovariectomized (Ovx) and one group was Sham operated (Sham). Eight weeks after surgeries, Sham-operated rats and one group of Ovx rats received control solutions of both PTH1-34 and Sr treatments. The three other groups of Ovx rats respectively received 625 mg/kg/day Sr (strontium ranelate; Servier Laboratory, Orléans, France) alone by oral gavage 5 days per week, 8 µg/kg/day PTH1-34 (Calbiochem) alone by subcutaneous injections 5 days per week, or both treatments for 8 additional weeks (curative treatments). The doses of Sr and PTH1-34 and treatment regimen were selected based on previously published studies in Ovx rats demonstrating robust anabolic effects on bone [11, 15, 20]. The day before sacrifice, rats were placed in metabolic cages, and urine was collected over an 18-hour period. At the end of the treatment period, rats were sacrificed, and their bones isolated for micro-computed tomography (µCT) analyses and mechanical testing. Experimental units were animals. At least 12 rats per group were required to detect a difference of 6% in cortical thickness (SD=10%) between groups at the significance level of 0.01 and a power of 90%. All tests were performed by a technician blinded to the treatment of each group.

### μCT

Trabecular bone microarchitecture of L4 vertebral bodies (100 slices from the beginning of secondary spongiosa) and cortical bone geometry of tibial midshafts (50 slices) were analyzed using a high-resolution μCT system (μCT 40; Scanco Medical, Basserdorf, Switzerland) employing a 12-μm isotropic voxel size as previously described [11].

### Nano-indentation

A nano-hardness test system (CSM Instruments, Peseux, Switzerland) was used to evaluate the intrinsic mechanical properties of trabecular bone tissue by recording the force shifts of a pyramidal diamond indenter pressed into the bone [11]. The L4 vertebral body of each rat was dissected from the intervertebral discs, embedded in polymethylmethacrylate, and cut transversely through the middle as previously described [11]. Samples were rehydrated following a standardized protocol for 16 hours in saline solution. The mechanical tests included five indentations performed on the same trabecular node located at the posterior end of each vertebral body. This nodal region, which may involve two interconnected trabeculae, was selected to minimize local structural heterogeneity and to assess intrinsic bone tissue material properties. The mean value of the five indentations was used for each specimen. Indents were set to a 900-nm depth with an approximate strain rate of ɛ = 0.066 s^−1^ for both loading and unloading. At maximum load, a 5-second holding period was applied. The limit of the maximal allowable thermal drift was set to 0.1 nm/s. The load displacement curve obtained during indentation permitted the calculation of tissue hardness (the average pressure that the material can resist), elastic modulus (stiffness), and working energy (area under the curve).

### Bone mechanical properties

The lumbar spines and tibiae, which were excised and frozen immediately after sacrifices, were thawed at 7 °C overnight and warmed to room temperature before mechanical testing. The posterior pedicle arches were removed from isolated L5 vertebral bodies without damaging the cortical shell. The superior and inferior endplates were embedded in methyl-methacrylate (Technovit 4701; Heraeus Kulzer, Wehrheim, Germany) over approximately 1 mm on each side to obtain two parallel loading surfaces and ensure uniform axial compression. The resulting free vertebral body height subjected to compression was approximately 3 mm [20]. Tibial strength was evaluated by a three-point bending test performed at the mid-diaphysis with a support span length of 20 mm. All specimens were tested in the same orientation [21]. The mechanical resistance to failure was tested using a servo-controlled electromechanical system (Instron 1114; Instron Corp., High Wycombe, UK). The displacement and load were simultaneously recorded. Maximal load (expressed in N), stiffness (slope of the linear part of the load/displacement curve, expressed in N/mm), and energy (total energy absorbed, area under the load/displacement curve, expressed in N.mm) were determined.

### Biological marker

Total urinary deoxypyridinoline (DPD) was calculated using a kit from Metra Biosystems (Mountain View, CA, USA) according to manufacturer’s instructions after acid hydrolysis of urine collected from rats.

### Primary osteoblast cultures

Primary osteoblasts were isolated from long bones of wildtype mice as previously described [6]. Tibiae and femurs were dissected, flushed out, washed, cut and digested in α-MEM (Amimed, Bioconcept) containing 10% FBS and 1 mg/mL collagenase II (Sigma) for 90 minutes at 37 °C. Digested bone chips were washed several times and incubated in α-MEM containing 10% FBS (Gibco) at 37 °C in a 5%-CO_2_/95% air humidified atmosphere for 6 days to allow osteoprogenitor migration from bone fragments. At that point, cells and bone chips were trypsinized (with trypsin/EDTA from Sigma) and passaged at a split ratio of 1:3. At the second passage, bone chips were removed. Medium was changed every 2-3 days. Osteoprogenitors at passages 3-4 were used for *in vitro* experiments. Confluent cultures were incubated in osteogenic medium containing α-MEM, 10% FBS, 0.05 mM L-ascorbate-2-phosphate (Sigma) and 10 mM β-glycerphosphate (AppliChem GmbH) to induce osteoblast differentiation for 4 days. Osteogenic medium was replaced, and cell cultures were pre-treated with 1 mM strontium solution consisting of a 100:1 molar ratio of Sr²⁺ derived from strontium chloride (Merck KGaA) and ranelate derived from strontium ranelate (Servier Laboratory), reflecting the circulating ratio of these two substances observed in patients treated with the standard dose of 2 g/day [18], for 1 hour and then stimulated with 10^−7^ M bovine PTH (1-34) (Calbiochem) at least 48 hours after osteogenic medium renewal for 3 days.

### RNA isolation and real-time PCR

Total RNA was extracted from primary osteoblast cultures using Tri Reagent^®^ (Molecular Research Center) and purified using a RNeasy Mini Kit (Qiagen). Single-stranded cDNA was synthesized from 2 µg of total RNA using a High-Capacity cDNA Archive Kit (Applied Biosystems) according to the manufacturer’s instructions. Real-time PCR was performed to measure the relative mRNA levels using the QuantStudio 5 Real-Time PCR System with SYBR Green Master Mix (Applied Biosystems). The primer sequences are described in Supplementary Table 1. Melting curve analyses performed at the completion of PCR amplifications revealed a single dissociation peak for each primer pair. The mean mRNA levels were calculated from triplicate analyses of each sample. Obtained mRNA level for a gene of interest was normalized to β2-microglobulin mRNA level in the same sample.

### Statistical analysis

All values are reported as mean ± SD. In vitro experiments were performed in triplicate and independently repeated 3 times. Effects of ovariectomy per se were analyzed by comparing Sham and Ovx rats using unpaired t-tests. Within Ovx rats, the effects of Sr and PTH1-34 treatments were analyzed using a 2 × 2 factorial design, and interactions between treatments were assessed by two-way ANOVA followed by Tukey’s multiple comparisons tests. Statistical analyses were performed by using the GraphPad Prism 10 software.

### Data availability

All data supporting the findings of this study are available from Zenodo (https://zenodo.org) with t}he identifier doi:10.5281/zenodo.15023891.

## Results

### Combined treatments with Sr and PTH1-34 further increased trabecular thickness and tissue-level volumetric bone mineral density in vertebral trabecular bone of Ovx rats

To test whether the combination of PTH1-34 and Sr treatments could exert greater beneficial effects on bone mass and strength than the two monotherapies against osteoporosis, we treated Ovx rats that lost bone for 8 weeks with Sr, PTH1-34 or both combined treatments for 8 additional weeks and measured their trabecular bone microarchitectures at L4 vertebrae. Ovx rats lost 28.3% of trabecular bone volume (p=0.0002 vs Sham) (Fig. 1). Sr treatment alone did not improve trabecular bone volume in Ovx rats (Fig. 1), but significantly increased tissue-level (material) volumetric bone mineral density (vBMD) (+3.1% vs Veh, p<0.0001), an effect that was greater than that observed with PTH1-34 monotherapy (Fig. 1). In contrast, PTH1-34 treatment alone significantly increased trabecular bone volume (+107.6% vs Veh, p<0.0001) in Ovx animals by augmenting trabecular thickness (+55.3% vs Veh, p<0.0001) and number of trabeculae (+20.1% vs Veh, p<0.0001), and reducing trabecular separation (−23% vs Veh, p<0.0001) (Fig. 1). These architectural changes were accompanied by marked increases in apparent trabecular vBMD (+76.9% vs Veh, p<0.0001) and a significant, though more modest, increase in tissue-level vBMD (+2.4% vs Veh, p<0.0007) (Fig. 1). Although the combination of Sr and PTH1-34 treatments did not exhibit a superior anabolic effect on trabecular bone volume compared to PTH1-34 treatment alone (+5.9% vs PTH1-34 alone, p=0.5123), it induced a significantly greater increase in trabecular thickness (+9% vs PTH1-34 alone, p=0.0047) in Ovx rats (Fig. 1). This was associated with a trend toward a further increase in apparent trabecular vBMD compared with PTH1-34 monotherapy (+9.6%, p=0.053). Notably, the combined treatment resulted in a significantly greater increase in material vBMD compared with Sr alone (+5.1%, p<0.0001) (Fig. 1).

**Fig. 1.**
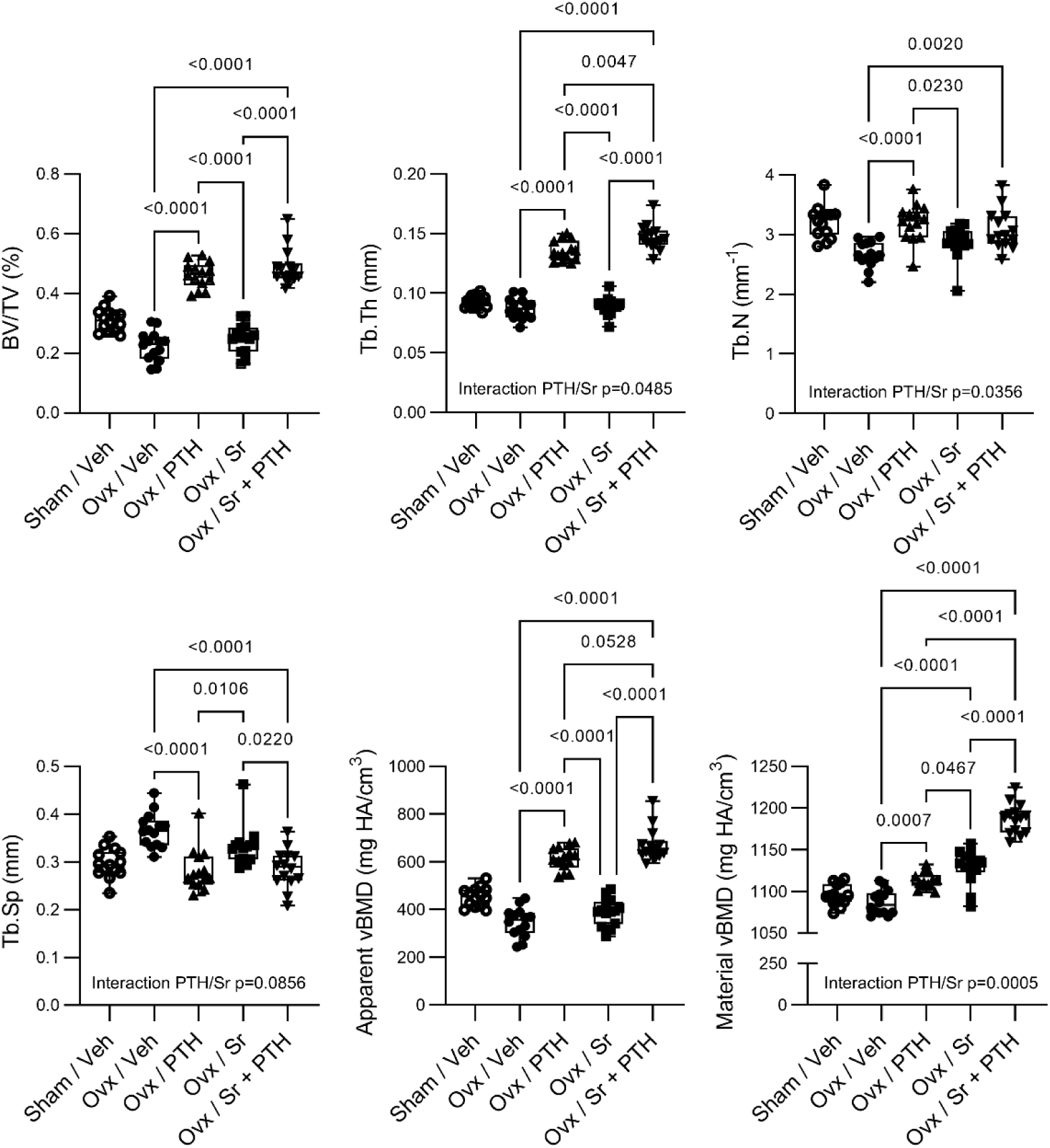
Sr treatment potentiates PTH1-34-induced trabecular thickness gain in Ovx rats. Female rats were either Sham-operated or Ovx at 6 months of age. 8 weeks after operations, Ovx rats received either vehicle solution, 625 mg/kg/day Sr (5 days per week), 8 µg/kg/day PTH1-34 (5 days per week), or both combined treatments for 8 weeks (curative treatment). Trabecular bone microarchitecture was measured at the 4^th^ lumbar vertebra (n=14 per group). µCT parameters include BV/TV: bone volume/total volume, Tb.Th: trabecular thickness, Tb.N: trabecular number, Tb.Sp: trabecular separation, and vBMD: volumetric bone mineral density. Interactions between effects of Sr and those of PTH1-34 in Ovx rats were analyzed by two-way ANOVA, and comparisons between the different groups were analyzed by Tukey’s multiple comparisons tests.

### Combined treatments with Sr and PTH1-34 further increased intrinsic bone quality and strength in vertebral trabecular bone of Ovx rats

At the level of L4 vertebral trabecular bone, the nanoindentation tests revealed that estrogen deficiency did not cause significant alteration of the intrinsic tissue quality (Fig. 2). Moreover, treatments with Sr or PTH1-34 alone did not significantly change parameters of bone quality in Ovx rats (Fig. 2). In contrast, combined treatment with Sr and PTH1-34 significantly enhanced the force (+11.4% vs Veh, p=0.0371), the average tissue hardness (+9.3% vs Veh, p=0.3349; +19% vs PTH1-34 alone, p=0.0133), and the working energy (+14.5% vs Veh, p=0.0086) of vertebral trabecular bone in Ovx rats (Fig. 2).

**Fig. 2.**
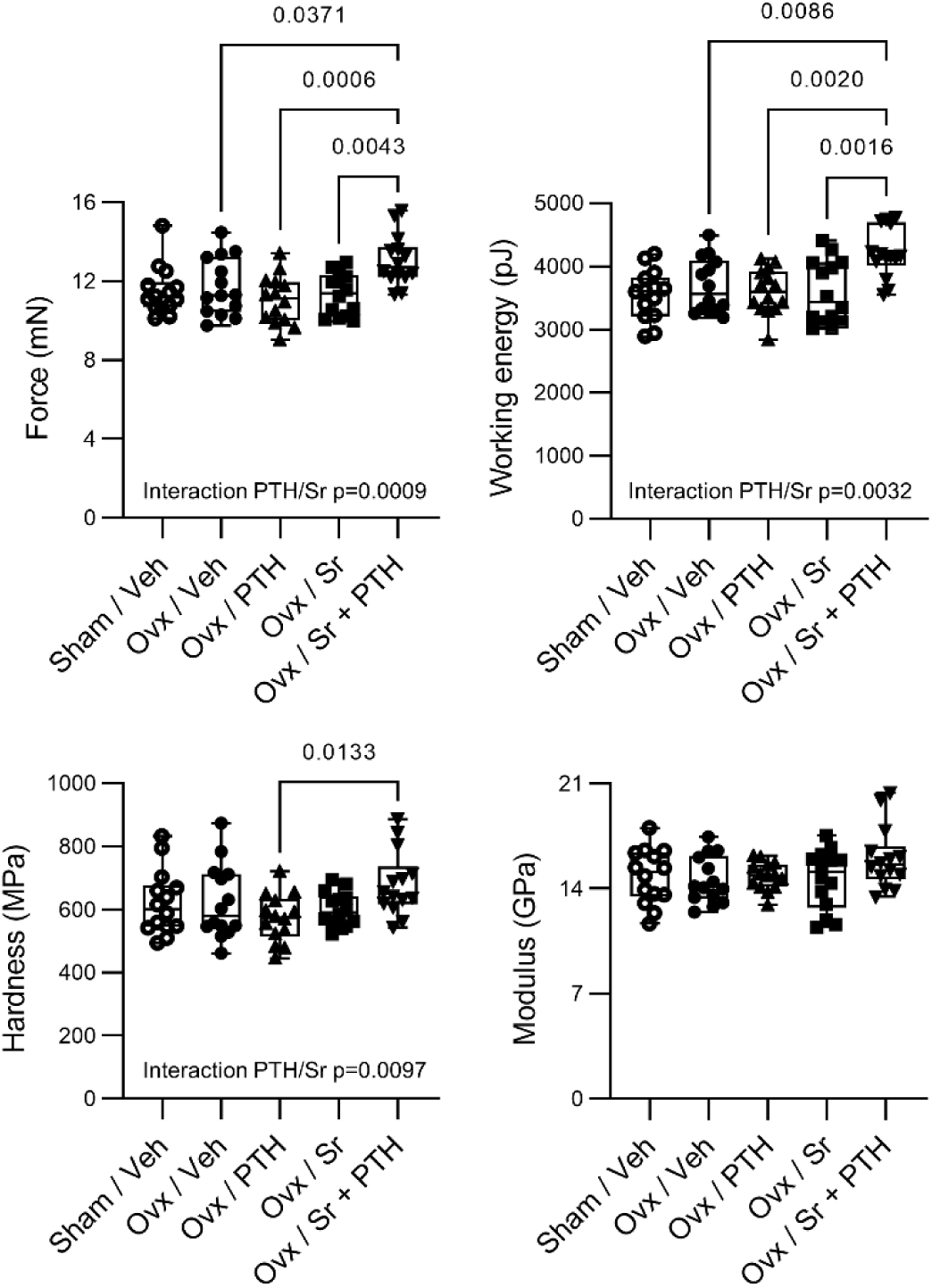
Sr treatment potentiates PTH1-34-induced increase in intrinsic bone tissue quality in Ovx rats. Female rats were either Sham-operated or Ovx at 6 months of age. 8 weeks after operations, Ovx rats received either vehicle solution, 625 mg/kg/day Sr (5 days per week), 8 µg/kg/day PTH1-34 (5 days per week), or both combined treatments for 8 weeks (curative treatment). Intrinsic bone tissue quality was assessed in trabecular bone at the 4^th^ lumbar vertebra by nanoindentation (n=14 per group). Interactions between effects of Sr and those of PTH1-34 in Ovx rats were analyzed by two-way ANOVA, and comparisons between the different groups were analyzed by Tukey’s multiple comparisons tests.

To explore whether the further increments of trabecular thickness and bone material properties induced by the combined treatment with Sr and PTH1-34 could be eventually translated into superior improvements in bone strength, we performed compressive tests on L5 vertebrae. The deleterious effects of Ovx were confirmed by significant decreases in maximal load (−14.5% vs Sham, p=0.0471) and total energy (−31.2% vs Sham, p=0.0026) (Fig. 3). The treatment with Sr did not show significant effect on biomechanical parameters of L5 vertebrae of Ovx rats (Fig. 3). The PTH1-34 treatment induced improvements of maximal load (+72.6% vs Veh, p<0.0001), total energy (+141.3% vs Veh, p=0.0003) and stiffness (+28.4% vs Veh, p=0.0899) in Ovx rats (Fig. 3). Interestingly, the combination of Sr and PTH1-34 treatments promoted further increases in maximal load (+26.3% vs PTH1-34 alone, p=0.0045) and total energy (+48.6% vs PTH1-34 alone, p=0.0033) in comparison to PTH1-34 treatment alone, and significantly augmented bone stiffness (+42.7% vs Veh, p=0.0038) in Ovx rats (Fig. 3).

**Fig. 3.**
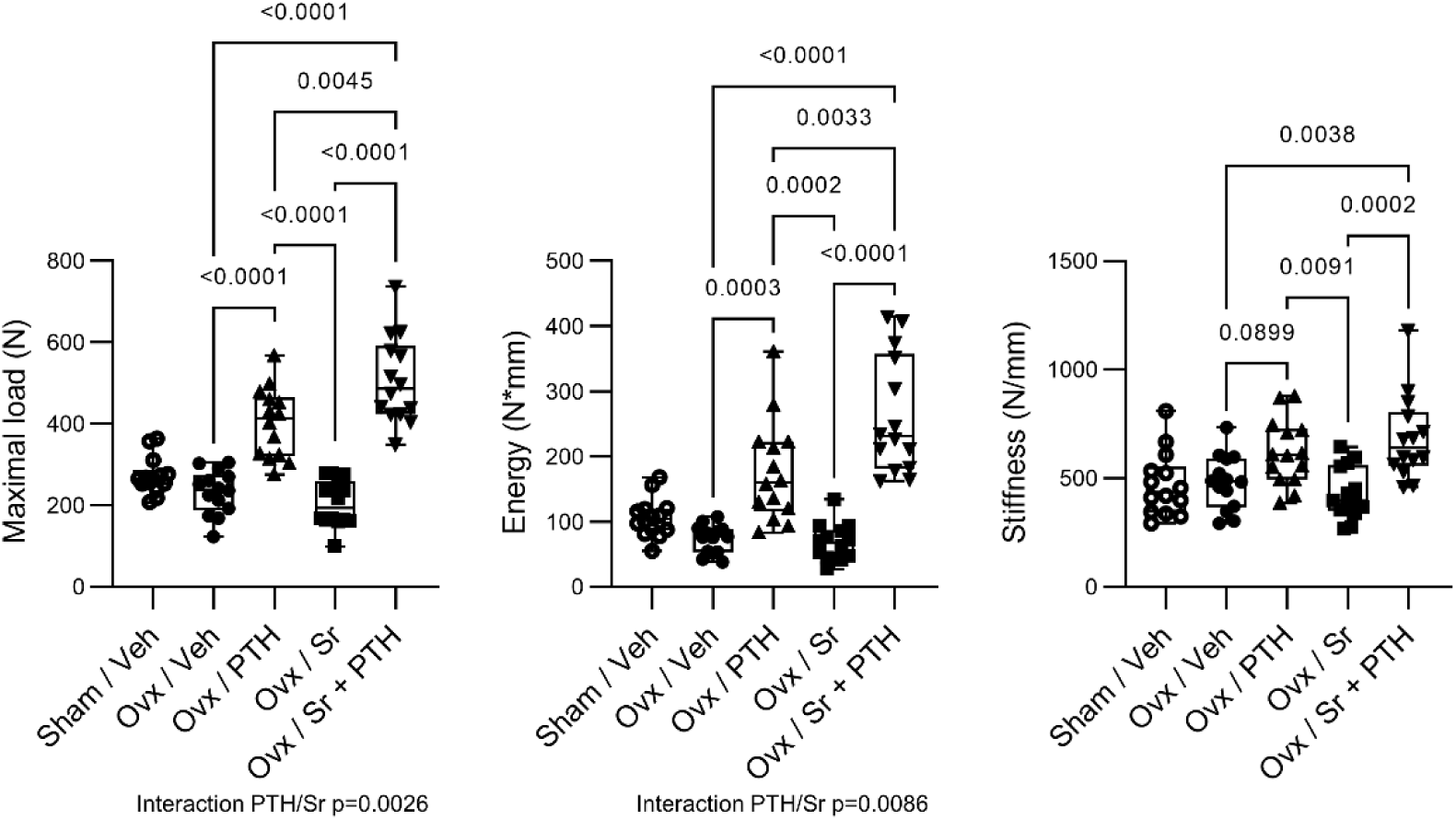
Sr treatment potentiates PTH1-34-induced increase in bone strength at lumbar spine of Ovx rats. Female rats were either Sham-operated or Ovx at 6 months of age. 8 weeks after operations, Ovx rats received either vehicle solution, 625 mg/kg/day Sr (5 days per week), 8 µg/kg/day PTH1-34 (5 days per week), or both combined treatments for 8 weeks (curative treatment). Bone mechanical properties were analyzed at the 4^th^ lumbar vertebra by axial compression tests (n=14 per group). Interactions between effects of Sr and those of PTH1-34 in Ovx rats were analyzed by two-way ANOVA, and comparisons between the different groups were analyzed by Tukey’s multiple comparisons tests.

### Only co-treatment with Sr and PTH1-34 significantly enhanced cortical thickness, apparent volumetric bone mineral density and strength in Ovx rats

The μCT analyses of tibial midshafts revealed a small reduction in cortical bone thickness in Ovx rats (−4%, p=0.1561) (Fig. 4). Sr monotherapy did not exert any beneficial effects on cortical bone geometry of Ovx rats, but significantly increased material vBMD (+2.8% vs Veh, p<0.0002) (Fig. 4). In contrast, the treatment with PTH1-34 significantly increased cortical bone volume (+13.6% vs Veh, p=0.0024) and thickness (+10.4% vs Veh, p=0.0052), but not cross-sectional area (unchanged Ct.TV) in Ovx animals (Fig. 4). This was associated with a significant increase in apparent cortical vBMD (+8.2% vs Veh, p<0.0001), but not tissue-level vBMD (Fig. 4). The combined therapy with Sr and PTH1-34 also enhanced cortical bone volume (+21.7% vs Veh, p<0.0001) and further augmented cortical thickness in comparison to PTH1-34 alone (+7.2% vs PTH1-34 alone, p=0.0558) (Fig. 4). This co-treatment also induced a significantly greater increase in apparent cortical vBMD compared with PTH1-34 monotherapy (+6.9% vs PTH1-34, p<0.0003), while tissue-level vBMD was not further increased compared with Sr alone (Fig. 4).

**Fig. 4.**
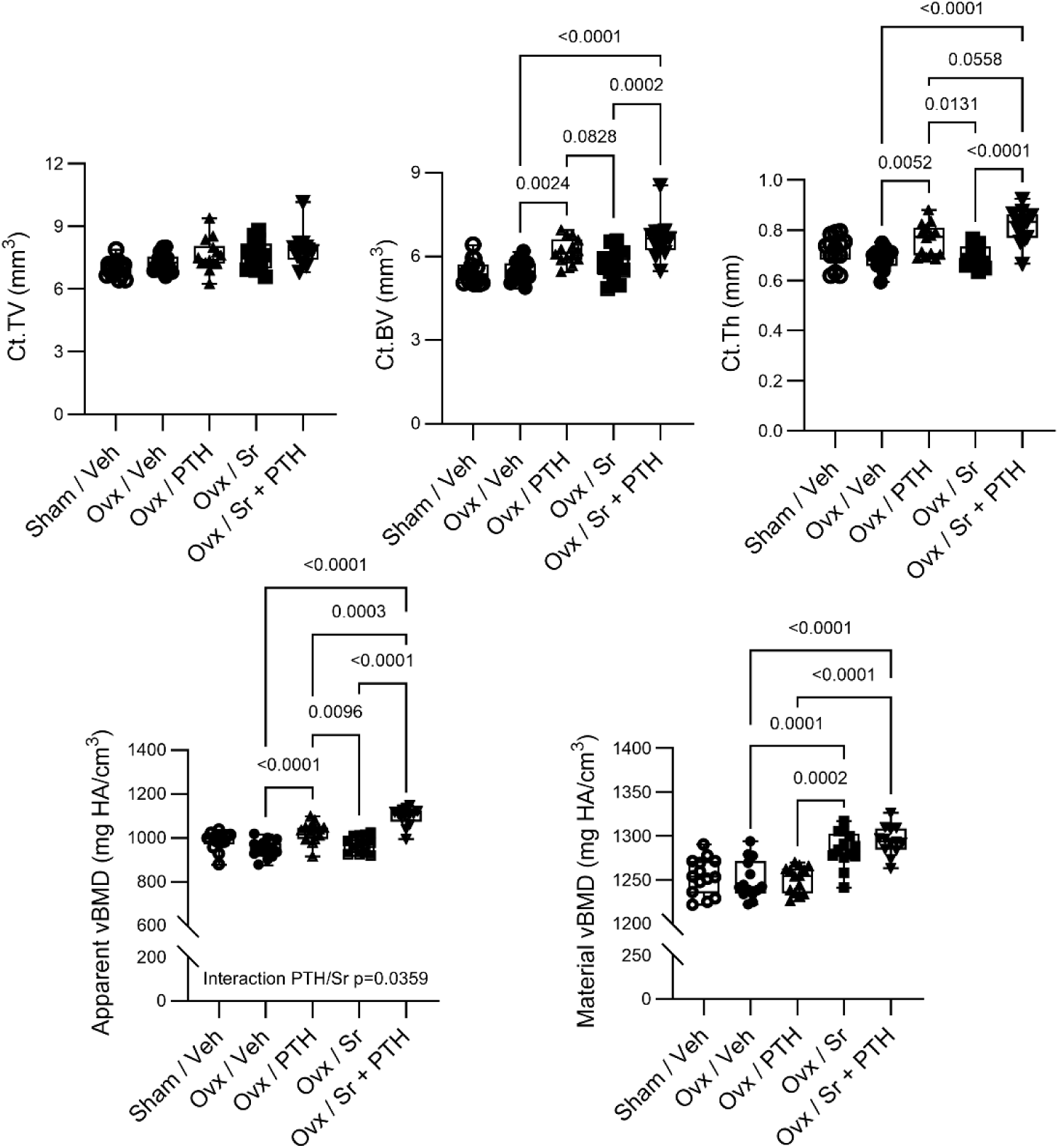
Combined treatment with Sr and PTH1-34 further enhanced cortical bone thickness in Ovx rats. Female rats were either Sham-operated or Ovx at 6 months of age. 8 weeks after operations, Ovx rats received either vehicle solution, 625 mg/kg/day Sr (5 days per week), 8 µg/kg/day PTH1-34 (5 days per week), or both combined treatments for 8 weeks (curative treatment). Cortical bone microarchitecture was measured at tibial midshaft (n=13 per group). µCT parameters include Ct.TV: cortical total volume, Ct.BV: cortical bone volume, Ct.Th: cortical thickness, and vBMD: volumetric bone mineral density. The two-way ANOVA did not detect significant interaction between effects of Sr and those of PTH1-34 in Ovx rats. Multiple comparisons were performed by using Tukey’s multiple comparisons tests.

The three-point bending tests performed in tibial midshafts did not show any changes in bone mechanical properties in Ovx animals (Fig. 5). Moreover, Sr or PTH1-34 monotherapies did not exhibit any effects on cortical bone mechanical properties (Fig. 5). Consistent with its superior effect on cortical thickness versus PTH1-34 treatment alone, the combined treatment with Sr and PTH1-34 exerted significant increases in maximal load (+20.7% vs Veh, p<0.0037) and working energy (+53.9% vs Veh, p<0.0025) (Fig. 5).

**Fig. 5.**
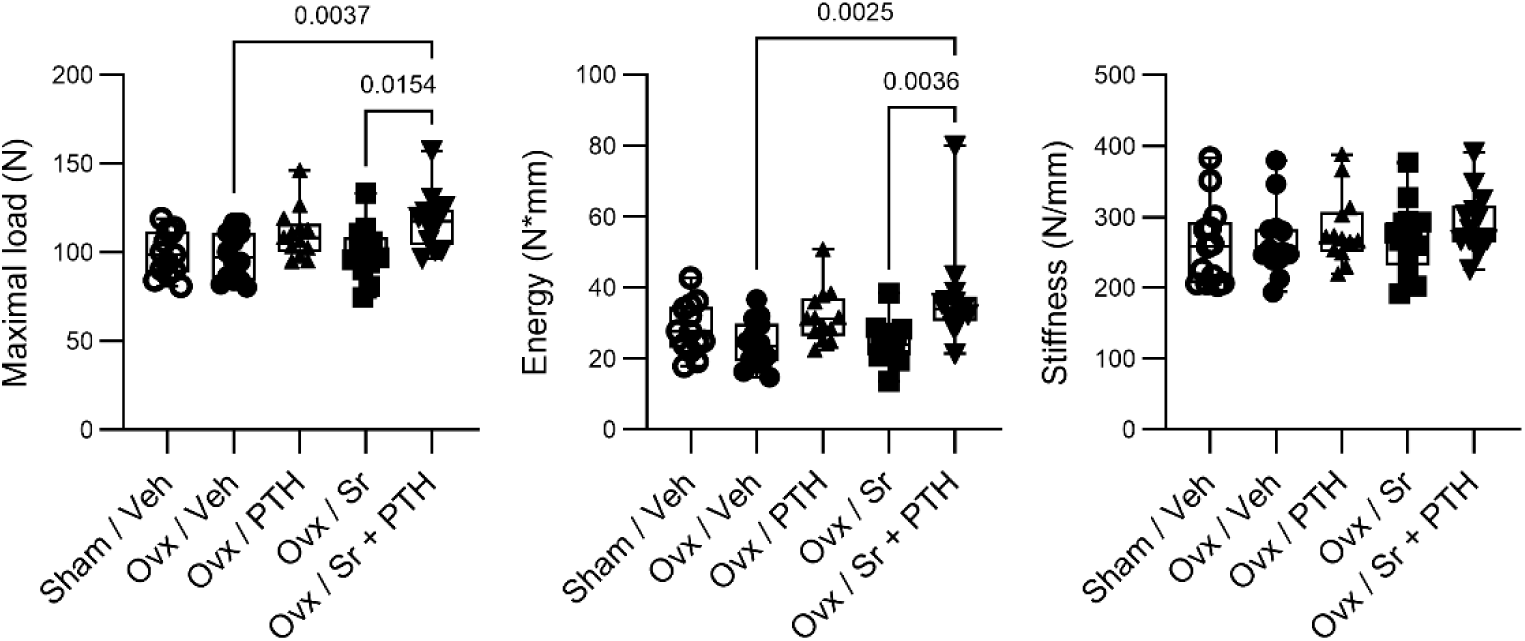
Only combined treatment with Sr and PTH1-34 exerts stimulatory effects on cortical bone strength in Ovx rats. Female rats were either Sham-operated or Ovx at 6 months of age. 8 weeks after operations, Ovx rats received either vehicle solution, 625 mg/kg/day Sr (Sr; 5 days per week), 8 µg/kg/day PTH1-34 (5 days per week), or both combined treatments for 8 weeks (curative treatment). Bone mechanical properties were analyzed at tibial midshaft by three-point bending tests (n=13 per group). The two-way ANOVA did not detect significant interaction between effects of Sr and those of PTH1-34 in Ovx rats. Multiple comparisons were performed by using Tukey’s multiple comparisons tests.

### The combined treatment with Sr and PTH1-34 increased bone resorption, but also further stimulated expressions of osteoblast differentiation markers in vitro

The effects of Sr and PTH1-34 monotherapies and co-treatment on bone resorption were assessed by measuring urinary levels of DPD. The urinary DPD levels were slightly and non-significantly increased in Ovx rats (Fig. 6). Although Sr treatment elevated urinary DPD levels in Ovx rats, this effect was not significant (+21.6% vs Veh, p=0.1203) (Fig. 6). Both PTH1-34 monotreatment (+29% vs Veh, p=0.0225) and Sr/PTH1-34 co-treatment significantly enhanced urinary DPD levels in Ovx rats (+36.4% vs Veh, p=0.0027), indicating that both treatments stimulated bone resorption in Ovx rats (Fig. 6).

**Fig. 6.**
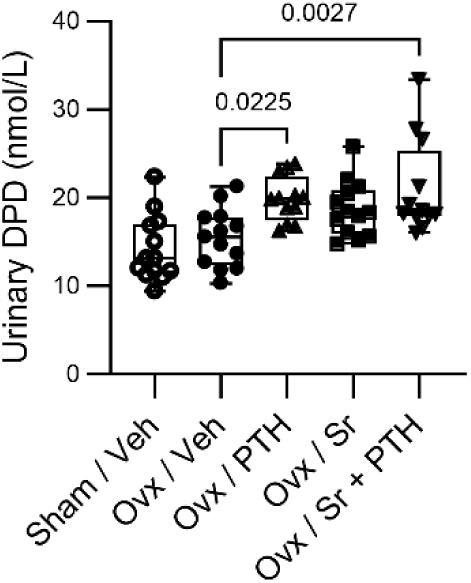
Sr, PTH1-34 and combined treatment increase bone resorption in Ovx rats. Female rats were either Sham-operated or Ovx at 6 months of age. 8 weeks after operations, Ovx rats received either vehicle solution, 625 mg/kg/day Sr (5 days per week), 8 µg/kg/day PTH1-34 (5 days per week), or both combined treatments for 8 weeks (curative treatment). Bone resorption was assessed by measuring urinary levels of deoxypyridinoline (n=12 per group). The two-way ANOVA did not detect significant interaction between effects of Sr and those of PTH1-34 in Ovx rats. Multiple comparisons were performed by using Tukey’s multiple comparisons tests.

To get further insights into the effects of Sr and PTH1-34 monotherapies and co-treatment on osteoblast differentiation and function, we treated primary cultures of murine osteoblasts with Sr, PTH1-34 or both treatments for 3 days. PTH1-34 treatment increased expression of *Rankl* and decreased that of *Opg* (encoding osteoprotegerin) (Fig. 7). While Sr treatment had no significant effect on *Rankl* expression, it also reduced that of *Opg* (Fig. 7). Stimulations with both Sr and PTH1-34 increased expression of *Rankl* and decreased that of *Opg* (Fig. 7). Those results were consistent with the higher bone resorption in Ovx rats receiving PTH1-34 alone or both Sr/PTH1-34 co-treatment (Fig. 6). Stimulations with Sr or PTH1-34 alone significantly increased expressions of *Igf1* (encoding insulin-like growth factor 1) and *Alpl* (alkaline phosphatase), but did not show any effects on those of *Col1a1* (type I collagen α1) and *Bglap* (osteocalcin) (Fig. 7). Interestingly, combined stimulation with Sr and PTH1-34 further enhanced expressions of *Igf1* and *Alpl*, consistent with a synergistic effect of Sr and PTH1-34 treatments on bone anabolism.

**Fig. 7.**
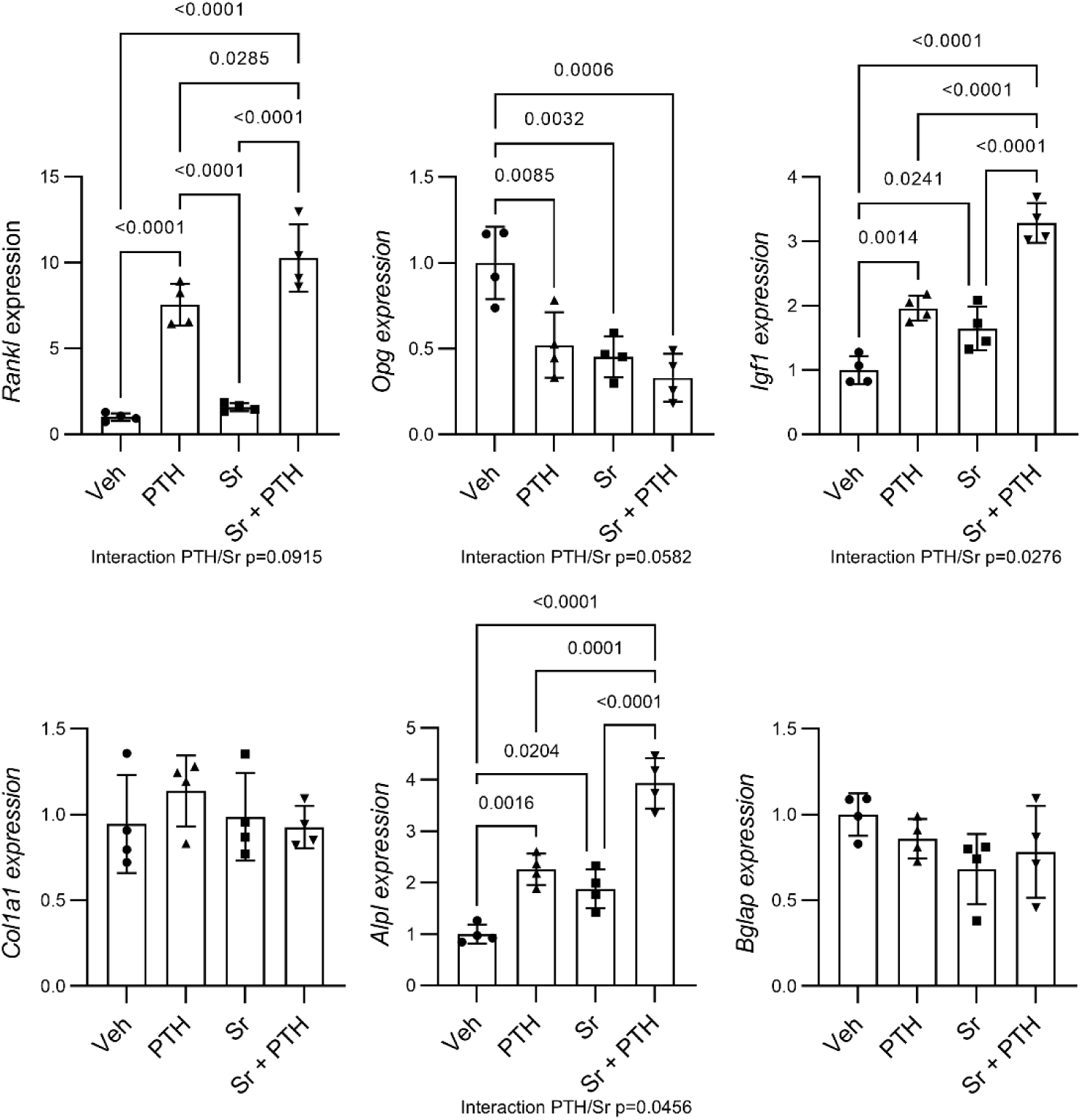
Sr treatment potentiates PTH1-34-induced expressions of receptor activator of NFκB ligand, insulin-like growth factor I and alkaline phosphatase in primary osteoblast cultures. Primary osteoblasts were isolated from mouse long bones, cultured in osteogenic medium, pre-treated with 0.5 mM strontium chloride (Sr) for 1 hour, and then stimulated with 10^−7^ M PTH1-34 for 3 days. Expressions of *Rankl* (encoding receptor activator of NFκB ligand), *Opg* (encoding osteoprotegerin), *Igf1* (encoding insulin-like growth factor I), *Col1a1* (encoding type I collagen α1), *Alpl* (encoding tissue non-specific alkaline phosphatase) and *Bglap* (encoding osteocalcin) were measured by quantitative RT-PCR (n=4 independent experiments). Interactions between effects of Sr and those of PTH1-34 were analyzed by two-way ANOVA and *post hoc* analyses were performed using Tukey’s multiple comparisons tests.

## Discussion

To optimize osteoporosis therapy with PTH1-34, we sought to determine whether Sr could potentiate the bone anabolic action of intermittent PTH1-34 in Ovx rats. We showed that co-treatment with Sr and PTH1-34 further increased trabecular thickness, apparent and tissue-level vBMD, bone material properties and trabecular bone strength in comparison to PTH1-34 treatment alone (Sr treatment alone only exerting a significant effect on tissue-level BMD) in Ovx rats. In addition, while PTH1-34 treatment enhanced cortical bone volume, cortical thickness and apparent cortical vBMD, co-treatment with Sr and PTH1-34 further elevated cortical thickness and apparent vBMD, while maintaining the increase in tissue-level cortical vBMD induced by Sr, and exerted a significant beneficial effect on cortical bone strength in Ovx rats. Co-treatment of primary osteoblast cultures with Sr and PTH1-34 increased *Rankl* expression and decreased that of *Opg*, which was consistent with elevated urinary deoxypyridinoline levels reflecting higher bone resorption in Ovx rats receiving the same treatment. Finally, co-stimulation with Sr and PTH1-34 further enhanced *Igf1* and *Alpl* expressions in osteoblast cultures, indicating a synergistic effect of both agents on bone anabolism.

The curative Sr monotherapy in Ovx rats did not exhibit any significant beneficial effects on bone volume, intrinsic properties, and strength at trabecular and cortical sites (Fig. 1-5). These differences with previous preclinical investigations were possibly due to the therapeutical approach employed here (curative vs preventive) and the relatively short period of treatment (8 weeks vs 16 to 52 weeks) [22, 23]. Nevertheless, Sr monotherapy consistently increased tissue-level vBMD at both trabecular and cortical sites (Fig. 1, 4), indicating a direct effect on bone mineralization rather than on bone mass or architecture. This effect is consistent with the ability of Sr²⁺ to substitute for Ca²⁺ within the bone mineral phase [19]. Surprisingly, Sr monotherapy exerted a non-significant elevation of bone resorption (Fig. 6), potentially due to a decreased *Opg* expression by osteoblast lineage cells (Fig. 7). This discrepancy with earlier studies having reported Sr anti-resorptive activity could provide a possible explanation for the absence of beneficial effect of our curative treatment strategy in Ovx rats [16, 17, 24]. Indeed, a short Sr treatment that started four weeks after Ovx in rats did not show any improvements in bone metabolism, mass, and strength [25].

The PTH1-34 monotherapy increased cortical and trabecular bone volumes, and improved trabecular bone microarchitecture in Ovx rats (Fig. 1, 4), leading to a marked increase in apparent vBMD, because of a rapid and strong anabolic effect reflected here by a stimulatory effect on osteoblast differentiation and *Igf1* expression (Fig. 7) [9, 10]. On the other hand, PTH1-34 failed to make an impact on material-level properties including tissue hardness and elastic modulus (Fig. 2). The absence of such an effect on intrinsic bone properties in response to PTH1-34 may reflect the poor collagen maturation and reduced mineralization degree due to accelerated bone remodeling (Fig. 6) [11]. Overall, those effects were translated into a marked increase in vertebral bone strength and a non-significant trend in elevated long bone strength (Fig. 3, 5), which is in agreement with the mechanisms of action by which intermittent PTH1-34 treatment reduces vertebral and non-vertebral fracture risk in osteoporotic patients [2, 3].

The concurrent therapy with Sr and PTH1-34 further augmented trabecular and cortical thicknesses (Fig. 1, 4), possibly through synergistic effects of Sr and PTH1-34 on osteoblast differentiation and *Igf1* expression as shown in vitro (Fig. 7). In this context, IGF1, which has been shown to increase in the serum of rats or osteoporotic women treated with Sr [15, 26], may represent a common signaling pathway required for the osteoanabolic action of both Sr and PTH1-34 [9, 10]. In addition to its effects on bone geometry, the combined treatment integrated the structural anabolic action of PTH1-34, reflected by increased apparent vBMD, with the mineral-level effects of Sr, which increased tissue-level vBMD and were maintained under co-treatment (Fig. 1, 4). Furthermore, unlike monotherapies, the combined treatment demonstrated beneficial effects on bone tissue properties such as tissue hardness and work-to-fracture (Fig. 2). The mechanisms underlying this response remain unclear, as bone remodeling was also stimulated by co-treatment with Sr and PTH1-34 (Fig. 6). However, it could be related to the combination of stimulated bone turnover and Sr ability to substitute for calcium within the bone mineral phase [19]. As a result, the combined therapy with Sr and PTH1-34 induced significantly greater improvements in vertebral and long bone strength in Ovx rats compared with monotherapies (Fig. 3, 5).

Interestingly, combination therapy with Sr and PTH1-34 appeared to produce notable skeletal benefits, including enhanced vertebral and non-vertebral bone strength, in rats with estrogen deficiency compared with previously reported combinations of PTH1-34 with antiresorptive agents [27–31]. Importantly, co-treatment with PTH1-34 and Sr improved bone quality (Fig. 2) and was associated with synergistic effects on markers of bone formation in vitro (Fig. 7), rather than merely expanding the anabolic window of PTH1-34 as seen with antiresorptive medications [27–31]. Although clinical trials of PTH1-34 combined with antiresorptive agents (such as estrogen, raloxifene, bisphosphonates, and denosumab) increased spine and hip bone mineral densities more than PTH1-34 or antiresorptive monotherapies, the additional benefit in fracture risk reduction was modest [32–34]. Based on our findings, it would be of interest to investigate the anti-fracture effectiveness of the concurrent therapy with PTH1-34 and Sr in osteoporotic patients and to explore whether this effect could be achieved with nutritional doses of Sr. Indeed, although strontium ranelate is no longer used as a therapeutic agent, Sr remains available as nutritional supplements and continues to be consumed for bone health. Beyond the specific compound, the present findings highlight the ability of Sr²⁺ to enhance PTH-induced improvements in bone quality and strength, supporting the broader concept that adjunctive strategies targeting bone material properties may improve the effectiveness of anabolic osteoporosis therapies.

## Statements and Declarations

### Author Contributions

CT: Conceptualization; investigation; methodology; data curation; formal analysis; funding acquisition; project administration; supervision; validation; writing-original draft; writing-review and editing. IB: Investigation; methodology; data curation, formal analysis; validation; writing-review and editing. PA: Conceptualization; funding acquisition; project administration; supervision; validation; writing-review and editing.

### Compliance with Ethical Standards

All performed animal experiments were in compliance with the guiding principles of the *Guide for the Care and Use of Laboratory Animals* (8^th^ edition) and approved by the Ethical Committee of the University of Geneva School of Medicine and the State of Geneva Veterinarian Office.

