## Supplementary Table 1 for "Strontium treatment potentiates bone anabolic action of intermittent PTH in ovariectomized rats"

**Supplementary Table 1**. List of primers used in real-time PCR.

| Gene | Primer | Sequence |
|---|---|---|
| *B2m* | F | CACTGACCGGCCTGTATGCT |
| | R | GTATGTTCGGCTTCCCATTCTC |
| *Rankl* | F | GCACACCTCACCATCAATGC |
| | R | AGCCTCGATCGTGGTACCAA |
| *Opg* | F | GACAACGTGTGTTCCGGAAA |
| | R | GGTAGGAACAGCAAACCTGAAGA |
| *Col1a1* | F | CTGGCCTTGGAGGAAACTTT |
| | R | GCACGGAAACTCCAGCTGAT |
| *Alpl* | F | AGATGGCCTGGATCTCATCAGT |
| | R | GTTCAGTGCGGTTCCAGACATA |
| *Bglap1* | F | GGAGGGCAATAAGGTAGTGAACAG |
| | R | CACAAGCAGGGTTAAGCTCACA |
| *Igf1* | F | GCTGGTGGATGCTCTTCAGTT |
| | R | CGAATGCTGGAGCCATAGC |
